# Iron-loading is a prominent feature of activated microglia in Alzheimer’s disease patients

**DOI:** 10.1101/2021.01.27.428379

**Authors:** Boyd Kenkhuis, Antonios Somarakis, Lorraine de Haan, Oleh Dzyubachyk, Marieke E IJsselsteijn, Noel FCC de Miranda, Boudewijn PF Lelieveldt, Jouke Dijkstra, Willeke MC van Roon-Mom, Thomas Höllt, Louise van der Weerd

**Author notes:** These authors contributed equally (Boyd Kenkhuis, Antonios Somarakis). Corresponding author, Boyd Kenkhuis, Department of Human Genetics, Leiden University Medical Center, Albinusdreef 2, 2333 ZA Leiden, The Netherlands.

## Abstract

Brain iron accumulation has been found to accelerate disease progression in Amyloid β-positive Alzheimer patients, though the mechanism is still unknown. Microglia have been identified as key-players in the disease pathogenesis, and are highly reactive cells responding to aberrations such as increased iron levels. Therefore, using histological methods, multispectral immunofluorescence and an automated in-house developed microglia segmentation and analysis pipeline, we studied the occurrence of iron-accumulating microglia and the effect on its activation state in human Alzheimer brains. We identified a subset of microglia with increased expression of the iron storage protein ferritin light chain (FTL), together with increased Iba1 expression, decreased TMEM119 and P2RY12 expression. This activated microglia subset represented iron-accumulating microglia and appeared morphologically dystrophic. Multispectral immunofluorescence allowed for spatial analysis of FTL^+^Iba1^+^-microglia, which were found to be the predominant Aβ-plaque infiltrating microglia. Finally, an increase of FTL^+^Iba1^+^-microglia was seen in patients with high Amyloid-β load and Tau load. These findings suggest iron to be taken up by microglia and to influence the functional phenotype of these cells, especially in conjunction with Aβ.

## 1. Introduction

Alzheimer’s disease is the most common cause of dementia, and is defined by the presence of amyloid-β (Aβ) plaques and tau tangles. In addition, the brain’s resident innate immune cells, microglia, have been found to be at the centre-stage of the disease, as most identified risk genes are predominantly or even exclusively expressed in microglia [1,2].

Not only can microglia modulate Alzheimer’s disease, but many transcriptomic studies showed microglia to undergo the most pronounced changes in response to pathology. In mice, a subset of responding microglia has been found to lose their homeostatic molecular signature and transition into a so-called ‘disease associated microglial state’ (DAM) [3]. In humans, a comparable yet disparate state coined the human Alzheimer microglia (HAM) has been identified [4]. Upregulated genes in these subsets do not only indicate loss of homeostatic function and increased pro-inflammatory activation, but also dysregulated iron-metabolism, manifested via upregulation of the *FTL-gene* and downregulation of *FTH1* and *SLC2A11* [4,5]. *FTL* encodes the ferritin light chain (FTL) protein, the component of the major iron-storage complex ferritin responsible for the long term storage of iron. These transcriptomic findings coincide with previously observed ferritin^+^ microglia in Alzheimer’s disease [6,7]. Though increased iron concentration likely plays a role, the exact link between the two has not yet been established.

Iron accumulation, irrespective of microglial activation, on the other hand, has been reported in disease-affected areas in Alzheimer’s disease, using both in-vivo and post-mortem human MRI [8]. Several MRI and histology studies found high correlations between iron accumulation and cortical Aβ and tau spreading [9–11]. Clinically, increased iron concentrations were shown to accelerate cognitive decline in Aβ-positive Alzheimer patients, indicative of a disease-modifying role for iron accumulation [12,13]. Again, how iron accelerates cognitive deterioration is poorly understood.

Therefore, in this study we aimed to research the possible link between iron accumulation and functionally activated microglia, and finally, its relation with Aβ-plaques. We performed a comprehensive investigation of iron-accumulating microglia, and first identified that the iron-storage protein FTL, specifically reflected increased iron accumulation in microglia. Secondly, by using multispectral immunofluorescence and an inhouse automated cell-analysis pipeline, we found FTL^+^ microglia to show significant activation, shown via both downregulation of homeostatic markers TMEM119 and P2RY12 and dystrophic morphology, and to predominantly infiltrate Aβ-plaques. This provides evidence for iron dysregulation as a prominent feature of activated microglia in Alzheimer’s disease in humans.

## 2. Methods

### 2.1 Tissue acquisition

Brain autopsy tissue of the middle temporal gyrus (MTG) of 12 Alzheimer patients and 9 age-matched controls was collected at the Leiden University Medical Center (LUMC), Netherlands Brain Bank (NBB) and the Normal Aging Brain collection Amsterdam (NABCA). Patients were included based on clinical presentation and diagnosis was confirmed by a neuropathologist. The neuropathologists also evaluates Braak stage, based on Gallyas and Tau immunohistochemistry (IHC), and Thal phase based on Congo Red and Amyloid Beta IHC, in eighteen standard regions, according to the latest international diagnostic criteria [14–16]. Patient demographics are reported in Supplementary Table 1. All material has been collected with written consent from the donors and the procedures have been approved by the Medical Ethical committee of the LUMC and the Amsterdam UMC.

### 2.2 Histology and immunohistochemistry

Formalin fixed paraffin embedded (FFPE) tissue was serially cut into ten 5-μm-thick and four 10-um-thick sections. Consecutive 10-μm-thick sections were used for histological detection of iron using an enhanced Perl’s stain and IHC detection of Ferritin Light Chain (FTL). 5-μm-thick sections were used for staining of the microglia multispectral immunofluorescence (mic-mIF) panel (Supplementary Table 2) to verify expression of FTL in microglia/macrophages (Iba1), look at the activation state of these cells (P2RY12/TMEM119) and study the interaction with Amyloid β-plaques (Aβ). Finally, of three subjects, 20-μm-thick sections were obtained for 3D confocal imaging. Step-by-step histological and IHC optimization protocols, together with the imaging parameters, are reported in the Supplementary Methods. A step-by-step mIF protocol and further analysis of the described histological, IHC and mIF staining will be described in the following sections.

### 2.3 Microglia multispectral immunofluorescence (mic-mIF) panel

One 5-μm-thick section of each subject was stained with the mic mIF panel with the following protocol. Sections were deparaffinized with 3× 5 min xylene, rinsed twice in 100% alcohol and subsequently washed with 100% ethanol for 5 min. Endogenous peroxidases were blocked for 20 min in 0.3% H_2_O_2_/methanol, after which the slides were rinsed with 70% and 50% alcohol. Heat induced antigen-retrieval was performed by cooking the slides for 10 min in pre-heated citrate (10 mM, pH=6.0) buffer for 10 minutes. After cooking, excess buffer was removed and slides were cooled for 60 min. Non-specific antibody binding sites were blocked with blocking buffer (0.1% BSA/PBS + 0.05% Tween) for 30 min. Firstly, slides were incubated with anti-TMEM119 (1:250, Sigma Aldrich) diluted in blocking buffer overnight at RT. Slides were washed thrice with PBS and incubated with Poly-HRP secondary antibody for 30 min. Slides were washed again and incubated with the appropriate Opal tertiary antibody (1:100 in amplification diluent, Perkin Elmer) for 60 min, which causes permanent binding of fluorophore to the antigen site. All subsequent steps are performed in the dark where possible. Finally, the slides are placed back in citrate buffer and cooked in the microwave for 15 min to wash the primary antibody off. The same steps are repeated for P2RY12 (1:2500, Sigma Aldrich). After binding of the two antibodies amplified with Opal, slides are incubated with a primary antibody mix with anti-FTL (1:100, Abcam), anti-Aβ (17-24) (1:250, Biolegend) and anti-Iba1 (1:20, Millipore) antibodies, diluted in blocking buffer, overnight at room temperature. The next day, after three washes with PBS, slides are incubated with a secondary antibody mix of G-a-rIgG A594, G-a-mIgG2b A647 and G-a-mIgG1 CF680 (1:200, ThermoFisher), diluted in 0.1% BSA/BPS. Finally, the slides are washed and incubated with 0.1 μg/mL DAPI (Sigma Aldrich) for 5 min, after which they are mounted with 30 uL Prolong diamond (ThermoFisher).

### 2.4 Post-mortem MRI acquisition and analysis

MRI data and T2*-w severity scores were obtained from a previous study by Bulk *et al*. [10], on the same tissue-blocks. In this study, tissue blocks were put in proton-free fluid (Fomblin LC08, Solvay), and scanned at room temperature on a 7T horizontal-bore Bruker MRI system equipped with a 23 mm receiver coil and Paravision 5.1 imaging software (Bruker Biospin, Ettlingen, Germany). A gradient echo scan was acquired with repetition time=75.0 ms, echo time=33.9 ms, flip angle=25° at 100 μm isotropic resolution with 20 signal averages. Subsequently, cortices were assessed for changes in MRI contrast following a pre-defined scoring system.

### 2.5 Iron-positive cell identification

Whole slide scans of the histochemical iron staining were exported from Philips Intellisite digital Pathology Solution platform (Philips, the Netherlands) and imported into ImageJ. RGB images were converted into 8-bit greyscale images. Subsequently, while blinded for diagnosis, for each subject an optimal threshold was set to include DAB-positive intracellular iron depositions, but exclude extracellular background signal. The cortex of the MTG was delineated and the number of positive cells was determined using the ImageJ particle analyser, with a size threshold of 4–100 pixels. Subject AD5 was excluded from this analysis, as iron-accumulating cells could not be distinguished due to high extracellular iron load.

### 2.6 Single cell segmentation

Identification of the different microglia types was based on the amount of expressed proteins over the segmented area (Supplementary Fig. 2a). Hence, accurate segmentation of the whole microglia cell area is of paramount importance for our method. Solutions currently available for microglia cell segmentation (Abdolhoseini *et al*., 2019 [14], Inform, PerkinElmer) typically fall short of capturing the whole microglia area (Supplementary Fig. 2b). These are focused on either capturing the skeleton of the cells, without properly identifying the cell boundaries (Supplementary Fig. 2c), or segmenting the microglia’s soma excluding their processes, which in the acquired 2D images are typically detached from the soma (Supplementary Fig. 2d). As a result, a novel segmentation algorithm for this type of data was developed.

Identification of the entire cytoplasmic area of microglia cells is error-prone, especially in regions close to Aβ-plaques, where microglia cells are densely packed. This problem was tackled by starting with the identification of the microglia’s soma. This part the of the microglia cells should overlap with its nucleus and shows high intensity values, making it easily discernible. Segmentation of microglia nuclei and somas was performed using a customized level-set-based cell segmentation method [18]. The main algorithm parameters are the weight for the energy terms minimizing the perimeter (ν) and the area (μ), which are empirically selected for each segmentation task. Larger parameter values corresponds to smoother segmentation results. For the microglia nucleus segmentation, the DNA component image was used as input, and level-set parameters were set to ν=2 and μ=3. Similarly, for the microglia soma segmentation the summation of the intensity values of the membrane (TMEM, PRY12, FTL, Iba) component images was utilized as input and level-set parameters were set to ν=2 and μ=3. In both cases, level-sets were initialized with regions obtained using the Otsu thresholding method [19] which is robust to intensity variation between images originating from the white and grey matter. Additionally, somas and nuclei with a total area smaller than 50 and 30 pixels, respectively, were removed.

For the extension of the obtained segmentation to the whole cytoplasmic area, the approach previously described for soma was repeated with less strong regularization (ν=2, μ=2). The result of this step was a finer segmentation capturing microglia areas that are less bright than the soma. Connected components overlapping with the previously identified somas were regarded as microglia cells, whereas not overlapping components were considered as possible detached processes. At this step, in case a blood vessel was identified in an image, the Li thresholding method [20] was chosen over Otsu for the initialization of the level-sets algorithm, as it is less sensitive to the high intensity pixels representing the vessel. Vessels were defined as components larger than 4000 pixels, after Otsu thresholding of the autofluorescent component image.

For correct identification of microglia cells in the proximity of Aβ-plaques, the watershed segmentation was applied specifically to those cells whose cytoplasmic area is shared among multiple microglia somas [21]. Aβ-plaque identification was performed employing a semi-supervised approach using Ilastik [22].

Finally, branches identified within a 10 pixel radius from the region corresponding to each identified microglia soma were identified as detached processes and assigned to the microglia cell.

A sample resulting segmentation of entire microglial cells is illustrated in Supplementary Fig. 2e. For the evaluation of our algorithm, 186 cells were manually segmented, in 7 images from different subjects and regions. Our proposed segmentation framework outperformed the available segmentation solutions correctly capturing 153 cells (Inform: 12 cells, Abdolhoseini *et al*., 2019: 49 cells compare Supplementary Fig. 2G), with false positive 33 cells (Supplementary Fig. 2H) and false negatives 40 cells (Supplementary Fig. 2I). Among the correctly identified cells, median Dice’s similarity index [23] of 0.8 was achieved (Supplementary Fig. 2f).

### 2.7 Cell phenotype identification

Superimposing the segmentation masks onto the component image of all membrane markers, four mean intensity values were extracted for each cell. Afterwards, intensity values were normalized imposing Z-score transformation. For the definition of the different microglia cell types Phenograph [24], an unsupervised clustering method, was utilized. For Phenograph 100 nearest neighbours along with the default parameters were selected, in order to avoid overclustering due to the limited amount of markers. Subsequently, for each Phenograph identified cluster the variability of the single-cell marker expression values was examined (Supplementary Fig. 3) through a violin plot [25] indicating the variation in each cluster, in parallel with their expression patterns as illustrated in the composite images.

### 2.8 Analysis of cellular phenotypes

The median expression value of each marker for each phenotype was illustrated with a heatmap. The similarities among the identified phenotypes were observed from a t-SNE [26] embedding using the same input as in Phenograph and the default parameters. The t-SNE embedding was coloured according to the cluster of each cell, its cohort or its individual marker expression values [27].

To explore the differences between the Alzheimer patients and controls regarding their phenotypes and their spatial relationship with the Aβ-plaques, an interactive, data-driven pipeline described by [28] was utilized. First, using a version of raincloud plots [28] the phenotypes that exist predominantly in each cohort are identified and consequently, their relative position regarding the Aβ-plaques using a visual query system are explored. For the exploration of the variability in each subject and the validation of our findings, a customized version of the motif glyphs described in our previous work [29] was employed.

### 2.9 Statistical analysis

Firstly, variables were inspected for being gaussian distributed. If normally-distributed, data plots represent the mean and the standard deviation. For not normally-distributed data, data plots show the median with the corresponding interquartile range. Comparison of two continuous variables was performed using a two-tailed unpaired Student’s independent t-test (normally-distributed) or a Mann-Whitney U test (not normally-distributed). Paired normally distributed data were analysed using a two-tailed paired Student’s t-test. Bonferroni post-hoc analysis was performed, and a significance level of *P* < 0.05 was used. The linear correlation between identified number of cells and different pathological hallmarks was assessed using the Pearson correlation coefficient. All statistical tests were performed using GraphPad Prism (Version 8.00, La Jolla, San Diego, CA, USA).

## 3. Results

### 3.1 FTL^+^-microglia reflect iron accumulating microglia in Alzheimer’s disease

An enhanced Perl’s staining for iron revealed an abundance of iron-positive cells in the cortex of the MTG in Alzheimer’s patients. On further inspection, iron-positive cells showed characteristic microglia morphology with a small soma and many thin processes (Fig. 1a) and quantification indicated a significant increase of iron-positive cells in Alzheimer patients compared to controls (*P* = 0.0024; Fig. 1b). Additionally, iron-positive cells appeared to cluster in groups, something that was not observed in control patients (Fig. 1a). All MTG tissue blocks have also previously been scanned using T2*-w MRI, sensitive for paramagnetic substances such as iron. MRI images were scored based on alterations in signal intensity reflecting overall parenchymal iron accumulation and focal iron depositions, and were published by Bulk *et al*. [10]. An increase of iron-positive microglia appeared to be only present in cases with the highest MRI severity score, indicating a significant increase of iron-positive microglia only to occur in subjects with a pronounced macroscopic iron-phenotype (Fig. 1c). Subsequently we studied the correspondence of iron accumulation with altered expression of the main iron-storage protein ferritin light chain (FTL), as FTL is known to be expressed in microglia and oligodendrocytes, whereas heavy chain ferritin is primarily expressed by neurons in Alzheimer tissue [30]. The Perl’s staining and the FTL staining showed a highly similar staining pattern, with focal clusters of cells representing microglia morphology (Fig. 1d). Thus, increased expression of the main iron-storage protein FTL appears to reflect iron accumulation in microglial cells.

**Fig. 1.**
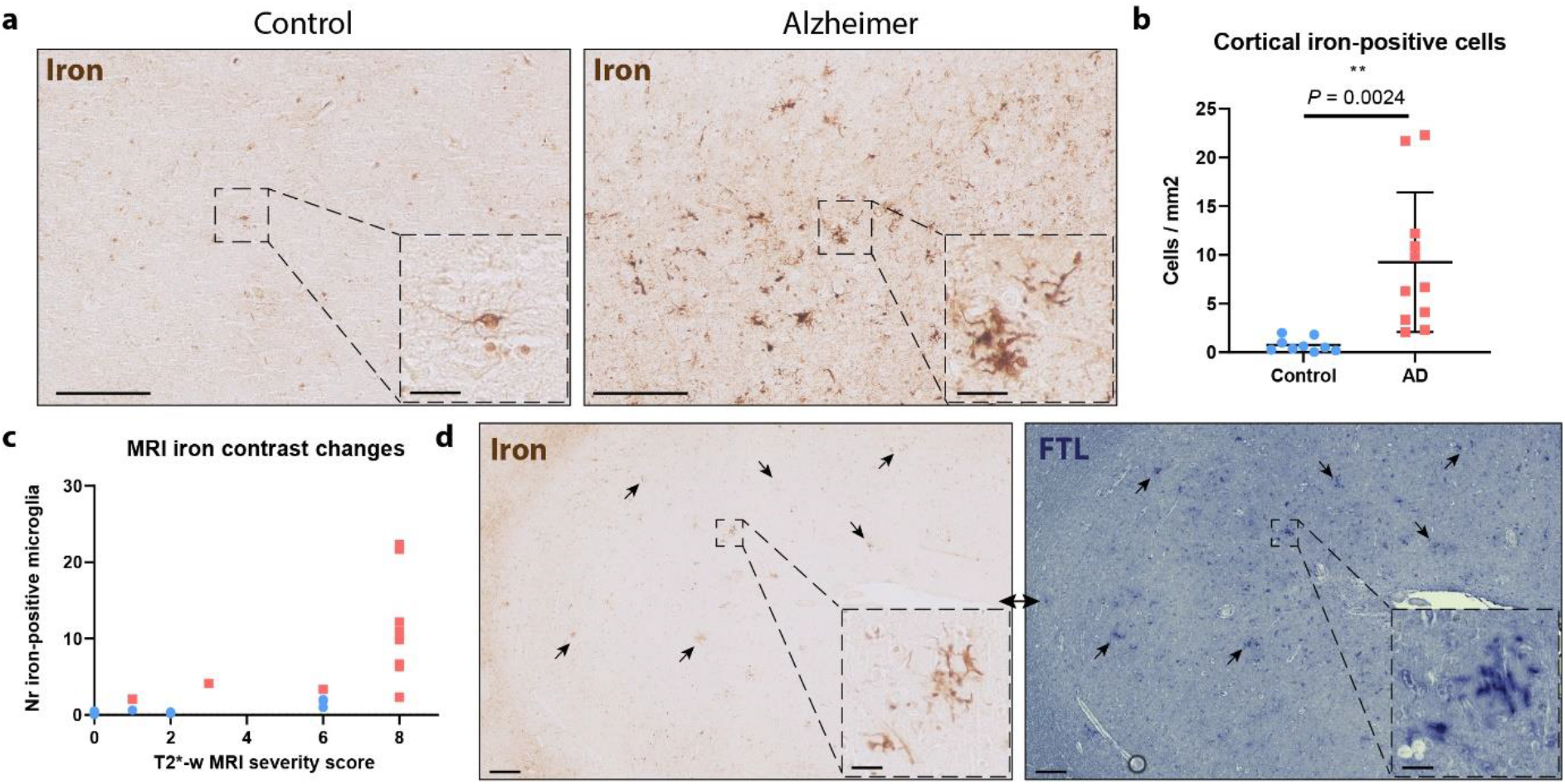
Increased iron-positive- and corresponding FTL^+^-microglia in Alzheimer’s disease. **a** MTG cortex of Alzheimer patients shows increased positivity for iron inside cells with microglial morphology. **b** Significant increase of iron-positive cells in Alzheimer patients (*n* = 11) compared to controls (*n* = 9)(Mean, Student’s t-test). **c** Iron-positive microglia number only increased in cases with severe signal alterations on iron-sensitive T2*-w MRI, reflected by MRI severity score. d FTL expression reflects intracellular iron accumulation. Scale overview images, 200 μm. Scale zooms, 30 μm.

### 3.2 Quantitative analysis enables microglia phenotyping

To confirm the microglial origin of FTL^+^ cells, study their activation state and potential interaction with Aβ, we designed the microglia multispectral immunofluorescence (mic-mIF) panel that can simultaneously detect 6 different markers (Supplementary Table 2). The MTG of 12 Alzheimer patients, both of early- and late onset, and 9 control subjects (Supplementary Table 1) was stained and imaged. After image acquisition and multispectral unmixing of the data, images were exported for automated segmentation, phenotyping and spatial analysis (Fig. 2). In total, 3149 images (110-236 per subject) were obtained. Multispectral unmixing allowed for simultaneous detection of FTL with the nuclear marker DAPI, TMEM119, P2RY12, Iba1 and Aβ at 0.5×0.5 μm resolution (Fig. 3a). TMEM119 and P2RY12 are generally considered homeostatic microglia-specific markers, based transcriptomic[3,31], *in vitro* [32–34] and post-mortem IHC studies [34–36], whose expression decreases when activated. Iba1, on the other hand, is a pan microglia/macrophage marker, which is upregulated upon activation. Finally Aβ stains the characteristic pathological Aβ-plaques that form in the parenchyma of Alzheimer patients. Images were segmented using a targeted in-house segmentation pipeline allowing segmentation of cells with processes (like microglia) in 2D images (Fig. 3b; Supplementary Fig. 2). After segmentation, unsupervised clustering using Phenograph assigned single segmented cells to 20 separate clusters. Following manual evaluation of the unsupervised clusters, 6 clusters were excluded based on non-microglial morphology and/or sub-threshold expression of all microglial markers (TMEM119/P2RY12/Iba1). In addition, three times two clusters were merged based on similarity in protein expression levels and their visual appearance (Supplementary Fig. 3). Exclusion of the non-microglial cells resulted in identification of 69227 cells, with no significant differences in the number of microglia per mm^2^ between control and Alzheimer patients in either grey matter (GM) or white matter (WM) (Fig. 3c). The remaining 11 clusters (C1-C11) were identified as major microglia phenotype clusters (Fig. 3d). Though the 11 different phenotypes clustered on the t-SNE plot, the low degree of separation suggests a rather continuous spectrum of expression of the microglia markers (Fig. 3e). The control and Alzheimer patients did cluster together, and the marker-based t-SNE plots already revealed more cells with high TMEM119 and P2RY12 expression in controls, but increased FTL expression in Alzheimer patients (Fig. 3f). With regard to anatomical region, only C1 and C2 appeared to be more present in the grey matter (GM), whereas C5 and C6 appeared to be proportionally more present in the white matter (WM) (Fig 3g). Four FTL^+^ clusters (C1–C3, C5) were identified, with differing expression levels and co-expression levels of P2RY12, TMEM119 and Iba1 (Fig. 3d). Cluster C1 (FTL^+^Iba1^+^) appeared significantly more present in Alzheimer patients (*P* = 0.0264), while C2 (P2RY12^+^TMEM119^+^FTL^+^Iba1^+^) was more present in controls (*P* = 0.0055; Fig. 3h). FTL^+^Iba1^+^ clusters lacking either P2RY12 (C3) or TMEM119 (C5) did not differ significantly in prevalence between control and Alzheimer patients. Cluster C4 showed solely Iba1 expression, meaning that this cluster likely also consists of non-resident infiltrating macrophages. Additionally, three P2RY12^+^ clusters (C6–C8) were identified, with the highest expressing cluster (C8) being more present in controls. The same applied for the TMEM119^+^ clusters (C9–C11), with C10 and C11 having higher expression and being more present in control patients. These results indicate a small shift of homeostatic microglia positive for P2RY12 and TMEM119 in controls towards activated microglia, with downregulated expression of P2RY12 and TMEM119 in Alzheimer patients. In addition, a specific Alzheimer-associated cluster shows increased expression of a combination of FTL and Iba1.

**Fig 2.**
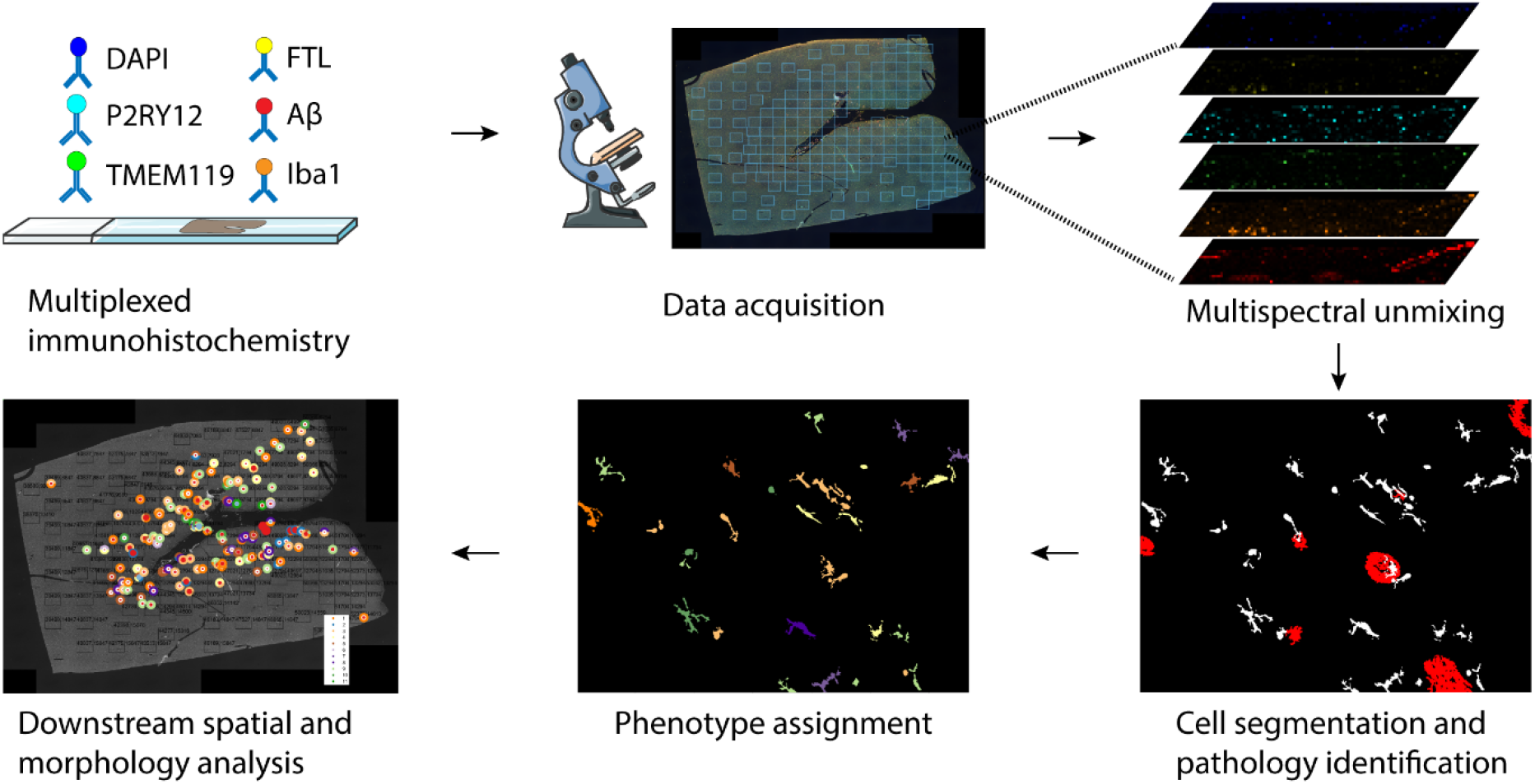
Schematic of mic-mIF acquisition and analysis pipeline.

### 3.3 Spatial analysis of FTL^+^ microglia clusters

After cell phenotype identification, all microglia were assessed for proximity to parenchymal Aβ-plaques. For visualization purposes, a second image was created, where infiltrated Aβ-plaques were plotted onto the original image as a ‘glyph’ (Fig 4a)[29], with the different colours corresponding to the respective cluster of the infiltrating microglia, to analyse which clusters predominantly infiltrated Aβ-plaques. Subsequently, all individual cells represented as cluster-colored dots or the cluster-colored glyphs were plotted back onto the original whole slide image (Fig 4a), to assess differences in cluster composition of microglial Aβ infiltration on a whole-section scale. As expected, quantification showed significantly more identified Aβ-plaques in Alzheimer patients, although some were found in controls as well (*P* = 0.0002; Fig 4b). Furthermore, a higher percentage of the plaques showed microglia infiltration in Alzheimer patients (*P* = 0.013; Fig 4c). Looking at the whole slide distribution, Aβ-plaques were found to be more present in the coronal sulcus rather than the gyrus. This also appeared to be associated with the regional microglia phenotype, as can be seen for the predominantly purple (C6–C8) microglia populating the Aβ-plaque deplete regions (Fig. 4d). To quantify the influence of Aβ-plaques on microglia phenotype, we compared all phenotyped microglia (all-mic) with the subset of microglia infiltrating Aβ-plaques (Aβ-mic). Controls showed a slight percental increase of C1 and C5 in Aβ-mic compared to all-mic, and less Aβ-plaque infiltration of TMEM119^+^ clusters C9–C11 (Fig. 4e), though this was based on a limited total number of Aβ-plaques. Alzheimer patients on the other hand, showed a large percental increase of FTL^+^-clusters C1 and C3 in the Aβ-mic population (Fig. 4E), which was also statistically significant when looking at subject-specific proportional increases (C1: *P* < 0.0001, C3: *P* = 0.0004; Figs. 4f,g). While C1 and C3 microglia together make up less than 20% of all-mic, they constitute almost 50% of the Aβ-mic population (Fig. 4e). P2RY12^+^ clusters C6–C8, on the other hand, showed a small contribution to Aβ-mic compared to all-mic (Fig. 4e). Finally, not only did C1 and C3 make up the majority of Aβ-mic, but also when examining the proportions of these individual clusters that directly infiltrated Aβ-plaques, they showed much higher proportion of infiltration than all the other clusters (Fig 4h). A visual example of the C1 and C3 microglia infiltrating an Aβ-plaque on the original mic-mIF images can be found in Fig. 4i. All in all, these results suggest Aβ-plaques to be predominantly infiltrated by a specific subset of microglia, characterized by increased FTL and Iba1 expression and loss of expression of homeostatic markers P2RY12 or TMEM119 and P2RY12.

**Fig 3.**
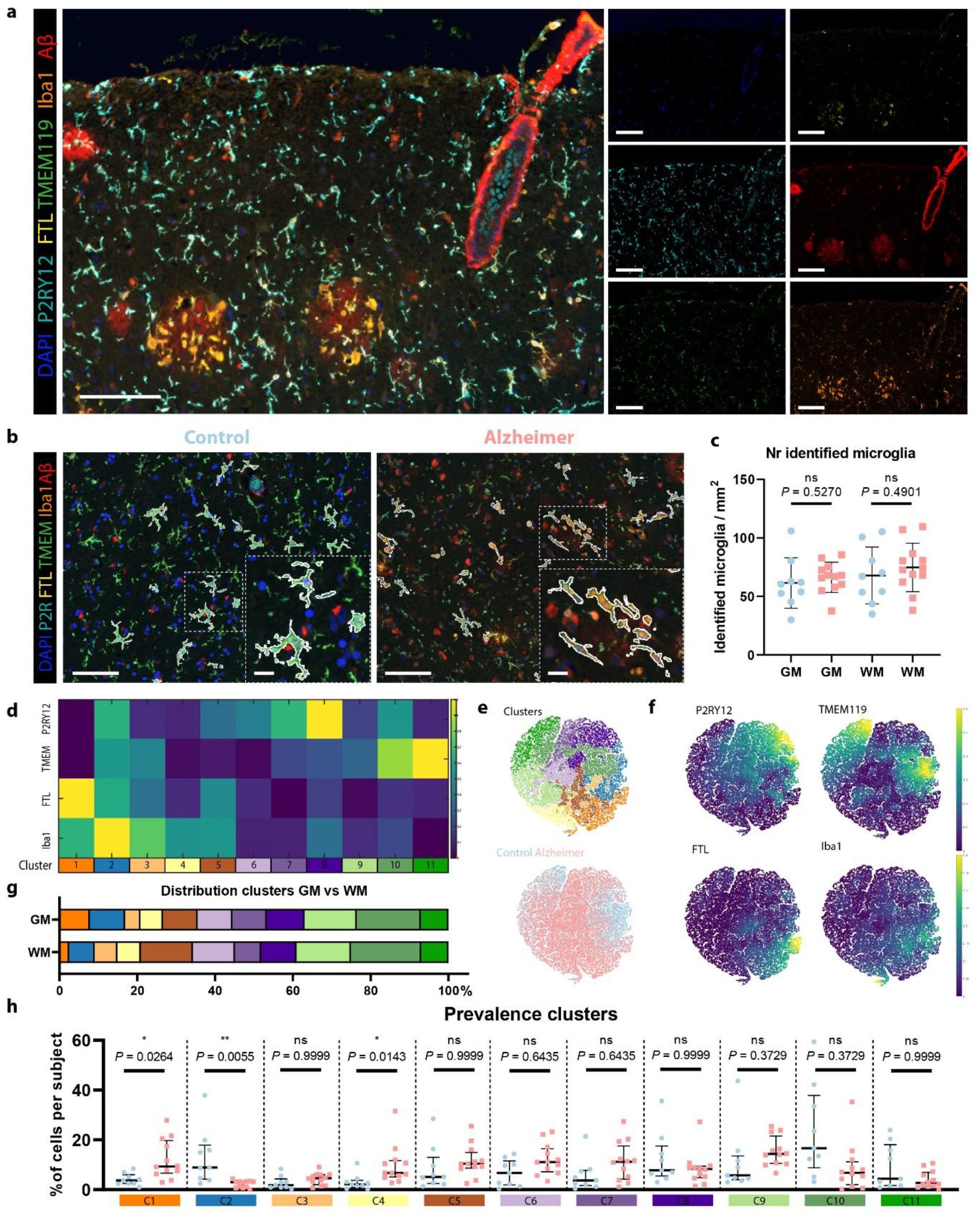
Identification of homeostatic and activated Alzheimer-associated microglia clusters. **a** Example of mIF image of an Alzheimer patient. **b** Exemplary images of segmented microglia in a control and an Alzheimer patient. **c** Number of identified cells in the GM and WM of controls (blue; *n* = 9) and Alzheimer patients (red; *n* = 12) (Mean, Student’s t-test). **d** Heatmap showing the expression of the four different markers (P2RY12, TMEM119, FTL and Iba1), in the 11 identified microglia clusters. **e** t-SNE plot of all individual cells showing the distinct colour-coded clusters and of control-vs. Alzheimer-patient-derived cells. **f** t-SNE plots colour-coded for intensity of the four individual markers. **g** Distribution of clusters in GM and WM. **h** Prevalence of identified clusters (C1–C11) in individual control (blue; n = 9) and Alzheimer patients (red; n = 12) (Median, Mann-Whitney U test). Scale bar, 100 μm. Scale bar zooms, 20 μm. GM = Grey matter, WM = White matter

**Fig 4.**
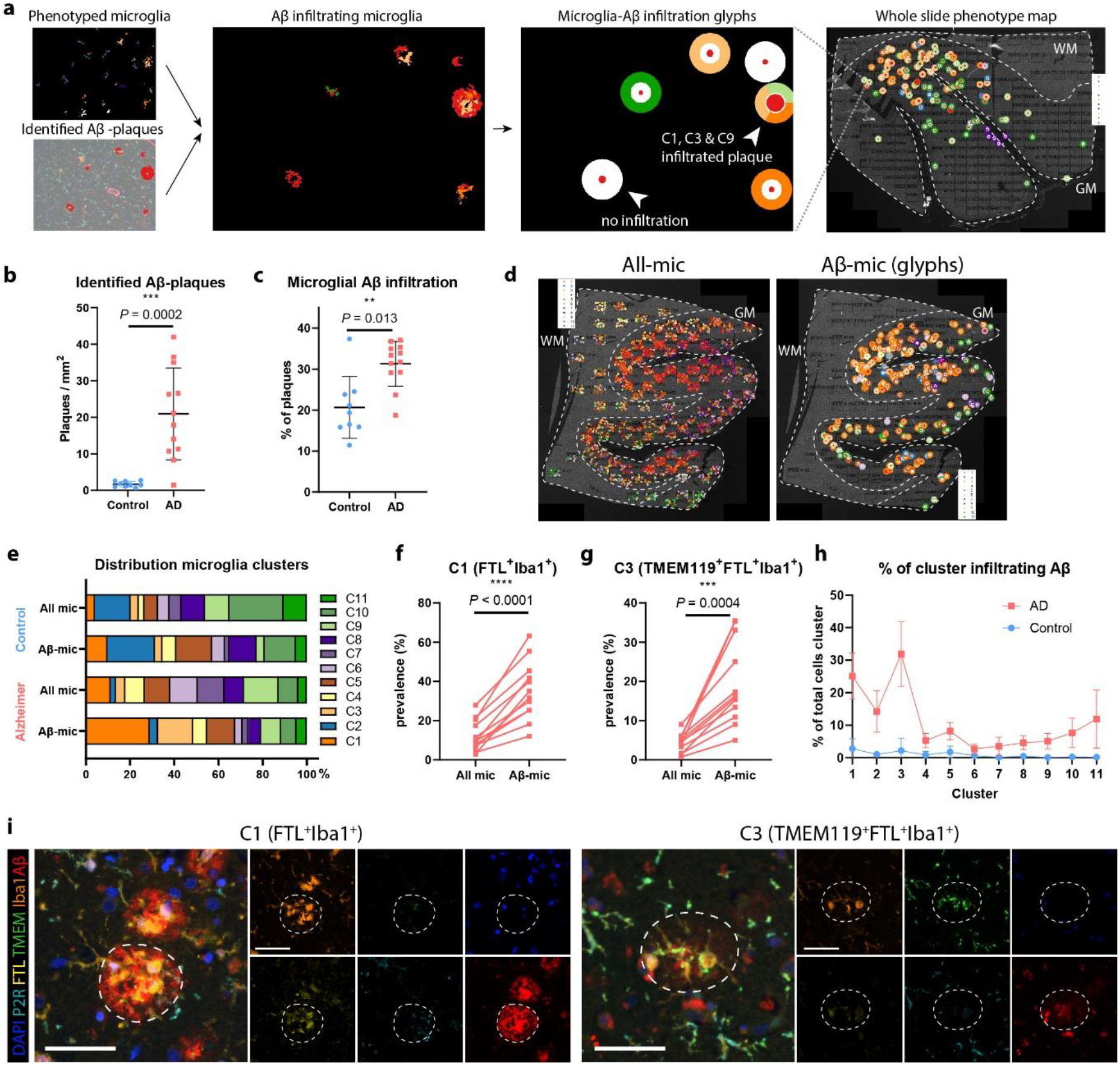
FTL^+^-microglia show significant Aβ-plaque infiltration. **a** Schematic of how microglial Aβ-plaque infiltration is studied. Both cells and Aβ-plaques are identified and an interaction map showing ‘glyphs’ in the colour of the cluster of the infiltrating microglia is created. Subsequently glyphs are plotted back onto the whole slide image to also enable studying the spatial distribution pattern. Number of identified Aβ-plaques (Mean, Student’s t-test) (**b**) and the percentage of microglia infiltrated Aβ-plaques (Mean, Student’s t-test) (c) are increased in Alzheimer’s disease (*n* = 12) compared to controls (*n* = 9). **d** Microglia clusters differ spatially, depending on the presence of Aβ-plaques in their proximity. **e** Distribution of all-mic clusters compared to Aβ-mic clusters of controls and Alzheimer patients. Comparison of prevalence of all-mic compared to Aβ-mic of C1-(**f**) and C3-microglia (**g**) of all individual Alzheimer patients (*n* = 12) (paired Student’s t-test). **h** Percentage of all identified clusters infiltrating Aβ-plaques. **i** Representative images of C1 and C3 microglia infiltrating an Aβ-plaque. Scale bar, 50 μm. All mic = all microglia, Aβ-mic = Aβ-plaque infiltrating microglia

### 3.4 Correlation of FTL^+^_microglia with pathology

As already shown in Fig. 1D, FTL staining closely followed the enhanced Perl’s staining showing microglial iron loading. Therefore, we also checked the correlation of the number of iron^-^positive microglia with the number of identified microglia of different FTL^+^ clusters. The number of identified C1 (FTL^+^Iba1^+^) microglia correlated well with number of iron--positive-cells (R = 0.7601, p = 0.0004; Fig. 5a), while other FTL^+^ clusters with lower expression (C2, C3) did not show correlation with number of iron-positive cells (Supplementary Figs. 4a,d)). This suggests that it is especially the marked increase of FTL expression found in C1-microglia that reflects substantial iron loading, while moderate FTL expression is also found in non-iron accumulating cells in controls. Although we already found C1-microglia to significantly infiltrate Aβ-plaques, we also checked for its correlation with overall Aβ and Tau load, as assessed by a neuropathologist using Thal stage and Braak stage, respectively. A marked increase of the number of C1-microglia was solely found in high-pathology load subjects with Thal phase V, and Braak stage V/VI (Fig. 5b,c), though not all high-pathology load subjects show increase of C1-microglia. C2-microglia were primarily found in controls with low Braak stage I/II and Thal I-II, whereas C3-microglia were present in both controls and Alzheimer patients with varying pathological burdens (supplementary Fig. 4b,c,e,f). This is in line with the finding that iron-positive microglia were particularly present in Alzheimer patients with advanced iron loading. However, there is lack of Alzheimer patients with intermediate Thal- and Braak-scores, making it impossible to state that an increase of C1-microglia is exclusive to advanced stage disease, and C3 represents an intermediate state between C2 in controls and C1 in advanced disease. Further investigation into the differences between early-onset Alzheimer’s disease (EOAD, onset <65y) patients and late-onset Alzheimer’s disease (LOAD, onset >65y) patients, showed no differences in Aβ load, microglia prevalence, or Aβ-infiltration of C1, C2 nor C3. (Supplementary Figs. 4g-q). In addition, we looked at differences between APOE3 and APOE4 carriers, as the latter have been found to have elevated ferritin levels in the CSF [37]. As expected, APOE4 carriers had more Aβ-plaques (Fig. 5d), but did not show overall increased microglia infiltration (Fig. 5e). Though sample sizes for both groups were small (*n* = 4-6), a trend indicating higher prevalence of C1-microglia in the GM could be observed (*P* = 0.0667; Fig. 5f), which was not the case for C2 and C3-microglia (Supplementary Fig. 4r,s) However, no difference was observed when looking at the proportion of Aβ-plaques infiltrated by C1 microglia (Aβ-mic) (*P* = 0.5096; Fig. 5g). This suggests that even though a higher percentage of C1-microglia infiltrate Aβ-plaques (*P* = 0.0381; Fig. 5h), this is likely due to the increased number of Aβ-plaques present in the APOE4 carriers.

**Fig 5.**
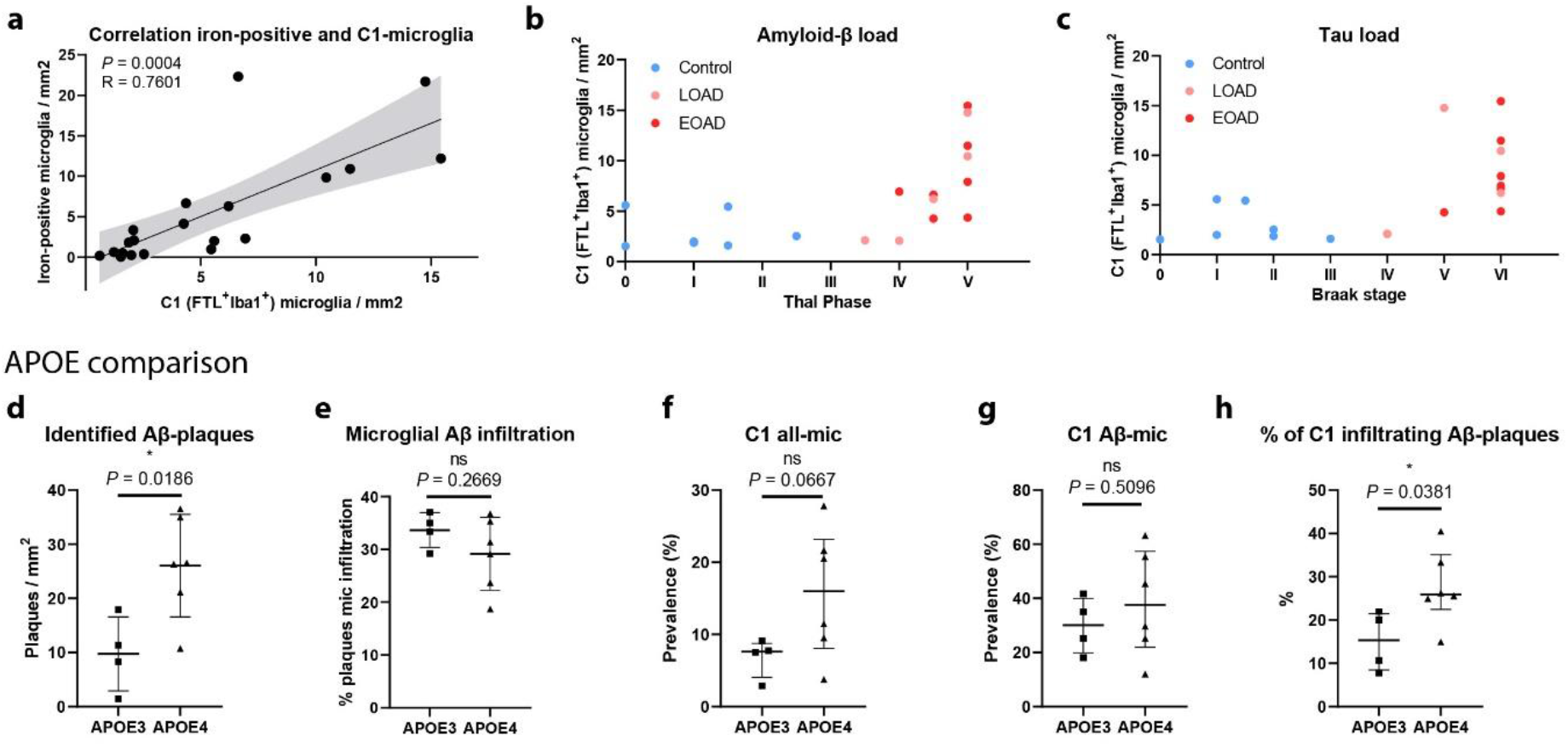
C1-microglia (FTL^+^Iba1^+^) reflect iron-positive microglia. **a** Number of identified C1-microglia correlates well with number of identified iron^+^-microglia (*n* = 20, Pearson coefficient). Increased number of C1-microglia are associated with higher overall Aβ load (Thal) (**b**) and Tau Load (Braak) (**c**). Comparison between APOE4 (*n* = 6) vs. APOE3 (*n* = 4) carriers shows increased number of identified Aβ-plaques (**d**) and similar microglia infiltration (**e**). Increased prevalence of C1-all-mic (**f**), no increased proportion of Aβ-plaques infiltrated with C1-mic (Aβ-mic) (**g**), and significantly increased proportion of C1-microglia infiltrating Aβ-plaques (**h**). **d-e** Median, Mann-Whitney U test. Patients AD8 and AD12 were excluded from the APOE comparison analysis as they harbour a familial mutation in the APP and PSEN1 gene, respectively, which could be of more influence than the APOE-genotype.

### 3.5 FTL^+^Iba1^+^-microglia have a dystrophic morphological appearance

Finally, we visually evaluated the morphological appearance of all phenotyped microglia in the same dataset, as this provides additional information about the activation stage of the microglia. Two authors (BK and LdH), evaluated the cells according to five distinctive morphological clusters: homeostatic, activated, dystrophic, phagocytic and perivascular macrophages (Fig. 6a), based on previously described morphological phenotypes [38]. The parenchyma of controls was predominantly populated by C6–C11-microglia, which consistently expressed TMEM119 and/or P2RY12. These cells presented with homeostatic morphology, showing small circular or oval cell bodies, with thin highly ramified processes and extensive branches (Fig. 6b). Morphological appearance therefore appeared to be in line with the homeostatic protein phenotype. Occasionally activated microglia were identified, which have larger cell bodies and noticeably fewer branches and ramifications (especially second degree) (Fig. 6a). Activated cells generally showed higher Iba1 and FTL expression and were often phenotyped as C2-microglia (Fig. 6a). Microglia in Alzheimer patients, on the other hand, had a much more heterogeneous appearance; homeostatic, activated, dystrophic and phagocytic microglia could all be observed within the coronal sulcus of a single patient (Fig. 6b). Though almost all phenotype clusters and morphological clusters could be observed, we focussed on the C1-microglia, as they reflected iron^+^-microglia. We found the striking majority of C1-microglia to have a dystrophic morphological appearance. The dystrophic cells show a very distinct phenotype, often with a cloudy or cytorrhexic (fragmentation of the cytoplasm) appearance which results in ill-defined processes (Fig. 6a). There is often deramification and the remaining branches show spheroids and fragmentation. Especially microglia (both C1 and C3) infiltrating Aβ plaques showed highly dystrophic morphological characteristics, indicative of an advanced activated/neurodegenerative state (Fig. 6c). The dystrophic morphology was also verified using 3D confocal microscopy, which also showed the same cytorrhexic appearance of microglia surrounding the Aβ-plaques (Fig. 6d). All in all, the finding of a dystrophic phenotype in C1-microglia was in line with the increased Iba1 and decreased TMEM119 and P2RY12 expression, which accompanied the pronounced FTL expression. They also reflected the morphological appearance of the iron-positive microglia identified on the Perl’s staining (Fig. 1a).

**Fig 6.**
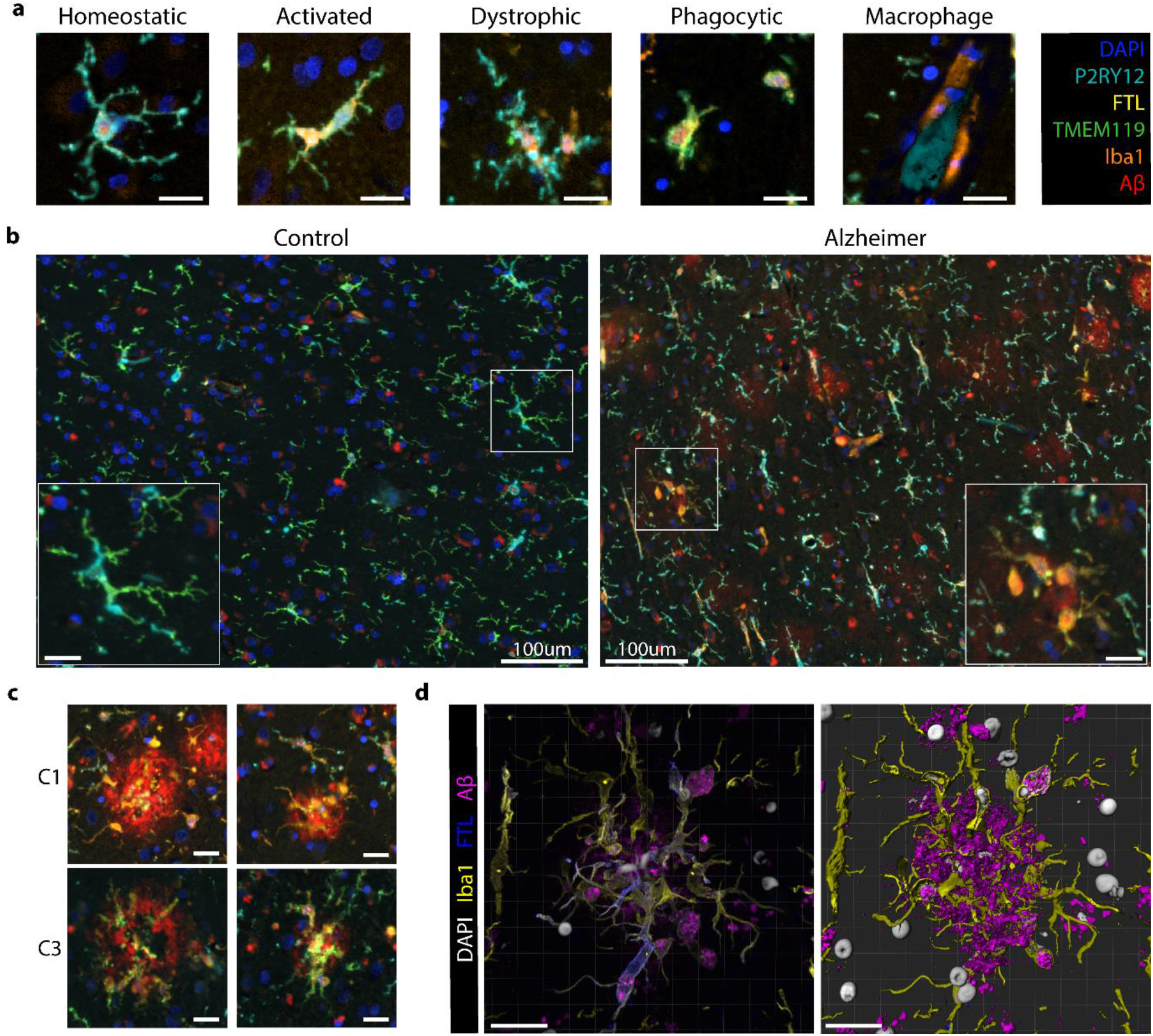
C1 and C3microglia show distinct dystrophic morphology compared to homeostatic control microglia. **a** Exemplary images of the five different morphological subtypes of microglia: homeostatic, activated, dystrophic, phagocytic and macrophage-like. **b** Controls show predominantly homeostatic and activated microglia, while Alzheimer patients show a variety of homeostatic, activated, phagocytic and dystrophic microglia. **c** Representative images of C1 and C3**-**microglia surrounding Aβ-plaques showing dystrophic morphology. **d** 3D confocal imaging confirms cytorrhexic appearance of FTL^+^Iba1^+^-microglia. Scale bar represents 20 μm unless otherwise stated. Colorcoding for IF-images in 6A-C are according to the box in the top right corner. Colorcoding of 3D confocal images are according to the legend adjacent to the images.

## 4. Discussion

In this manuscript, we confirmed that increased FTL expression reflects an increase in iron accumulation in microglia in the cortex of Alzheimer patients. Microglia with increased FTL expression also showed higher Iba1 expression, but loss of homeostatic markers TMEM119 and P2RY12, indicative of an activated phenotype. On further investigation this FTL^+^Iba1^+^ phenotype appeared to be increasingly present in Alzheimer patients and the predominant Aβ-plaque infiltrating microglia phenotype. Morphologically they appeared to be in a dystrophic activation stage.

Firstly, in this study we confirmed that previously identified iron-positive cells in Alzheimer patients [39,40] are of microglial rather than astrocytic origin, and show high FTL expression. Subsequently, using multispectral fluorescence and unsupervised clustering, we identified an several FTL^+^-clusters, which were variably present in controls and Alzheimer disease stages. C2-microglia, which displayed positivity for all included microglia markers were almost exclusively present in control patients. Conversely, C1-microglia (FTL^+^Iba1^+^) were significantly more present in AD patients, and C3-microglia (TMEM119^+^FTL^+^Iba1^+^) were marginally present in either group. Interestingly, both C1 and C3-microglia showed a strong tendency to infiltrate Aβ-plaques. C1-microglia were almost exclusively present in advanced stage Alzheimer patients, whereas C2-microglia were primarily detected in controls (with low Thal/Braak stages), and C3-microglia were variably present across controls and Alzheimer patients of all stages. Regarding the temporal dynamics of these clusters, one could therefore hypothesize that in Alzheimer’s disease microglia surround Aβ-plaques and lose P2RY12 expression, as has been observed previously by others (transition from C2 to C3) [35,41]. As of yet we do not know what the relevance is of the preserved TMEM119-expression. Over time, these microglia take up iron, causing a pronounced increase of FTL expression and loss of TMEM119. This corresponds to the fact that only C1-microglia appeared to correlate with iron-accumulating microglia. However, our study population is not ideal to dissect the temporal dynamics of these clusters, since the majority of Alzheimer patients showed advanced disease (Braak V/VI) and only two patients showed mild to moderate (Braak III/IV). Future work studying these phenotypes in a larger cohort with a larger range of disease stages would be highly relevant to accurately determine at what stage of the disease C2 microglia prevalence decreases and C1 and C3 microglia prevalence increases.

Several qualitative studies had previously identified increased presence of dystrophic ferritin^+^ microglia in brain tissue of Alzheimer patients [6,39,42,43]. The dystrophic morphological appearance was also confirmed in this study, though the functional insights of these morphologically defined states remains debatable. Our spatial analysis revealed a strong tendency of FTL^+^Iba1^+^ to infiltrate Aβ-plaques; significantly more than can be expected based on prevalence of the cluster itself, and more than any other identified microglia cluster. Although some other studies had already looked into the association of dystrophic ferritin^+^ microglia with Aβ-plaques [6,7,30,39,44], results were inconsistent, as none of these studies so far looked into the relative proportion of these microglia in the total population. The importance of this is also stressed in a recent study by Nguyen *et al*. [45], in which they found an amyloid-responsive microglia (ARM) subset, characterized by CD163, but did not pick up on the Aβ-plaque-infiltrating properties of their identified ferritin^+^ microglia. Finally, we were able to further characterize iron^+^/FTL^+^-microglia by analyzing co-expression of several other microglia markers on a single cell level. This revealed that C1-microglia, with the highest FTL protein expression and increased Iba1 expression, showed complete loss of expression of homeostatic markers TMEM119 and P2RY12. Although we acknowledge that our FTL^+^Iba1^+^(P2RY12^-^TMEM119^-^)-microglia were only characterized using four protein-markers, which is only a fraction compared to the total amount of genes used to define specific transcriptomic states such as the DAM/HAM-states, we do want to highlight the similarities. The HAM/DAM-subsets showed FTL among the highest upregulated genes, with coinciding downregulation of TMEM119 and P2RY12 [3,4]. Additionally clustering around Aβ-plaques was also reported as a characteristic feature of DAM-microglia [3], as is observed for the identified FTL^+^Iba1^+^-microglia.

To date, the reason for the observed increase of FTL-expression remains disputed. With FTL being the long-term storage component of ferritin, its expression is likely to be increased in response to increased intracellular labile iron concentrations. Yet, ferritin is also widely recognized as an acute phase reactant and it has also been suggested that microglia upregulate ferritin as a response to exhaustion, caused by the attempting to phagocytose aggregated Aβ [44]. However, our findings show that the identified FTL^+^Iba1^+^-microglia closely reflected microglia with high levels of the metal iron, and therefore suggest that the observed increased FTL-expression at least does not merely reflect inflammatory activation or exhaustion, but also increased iron levels. This is in line with a previous study, which found ferritin levels in the CSF to not be associated with an inflammatory response in Alzheimer patients and hypothesized ferritin levels to rather reflect changes in iron associated with tangle and plaque pathology [46].

Why iron increases with age and even more profoundly in neurodegenerative diseases is still largely unknown [8,47]. It is hypothesized to be caused by several factors including increased blood-brain barrier permeability and disorganization of the iron-dense myelin sheaths[48–50]. Alongside a general increase of iron in the parenchyma, iron was also shown to accumulate inside Aβ-plaques [50,51]. Therefore, a possible hypothesis for why iron is sequestered in microglia surrounding Aβ-plaques, could be that the iron is taken up as byproduct while attempting to phagocytose the Aβ aggregates. Conversely, considering we only found approximately 25% of iron-accumulating C1-microglia to infiltrate Aβ-plaques, iron is more likely sequestered using either DMT1 or Transferrin-receptors and stored inside FTL, in an attempt to mitigate the potentially toxic effects of free iron, which in its free form is suggested to partake in Fenton’s reaction to form hydroxyl radicals and cause toxic oxidate stress [49].when iron is taken up by microglia, it first becomes part of the labile iron pool, where it can produce reactive oxygen species damaging the mitochondria and other cell organelles [52]. Studies performed using peripheral tissue cells showed the non-CNS equivalent of microglia, macrophages, to respond to intracellular iron accumulation by also activating the NLRP3 inflammasome [53]. Accordingly, *in vitro* and *in vivo* studies have shown that exposure to a combination of iron and Aβ induces the production of cytokine IL-1β and a switch to glycolytic metabolism in microglia, both of which can be interpreted as NLRP3-inflammasome activation [54,55]. NLRP3-inflammasome activation in microglia was shown to be able to modify disease progression in two different Alzheimer mouse models [56,57]. Our data support the *in vitro* and mouse model evidence that iron and Aβ can act together to accelerate disease progression via microglial inflammasome activation, by showing that in human brain tissue of Alzheimer patients, microglia are exposed to a combination iron and Aβ. Finally, these findings are also in line with recent clinical studies, in which iron was found to act as a potential disease modifier by accelerating deterioration in Alzheimer patients with high Aβ load [12,13].

Thanks to the possibility to visualize up to six protein markers on the same section using mIF, we could better study the great heterogeneity in microglia phenotype and its spatial relationship with pathology. A limitation of mIF compared to other high-dimensional techniques such as single-cell or imaging mass cytometry is the limited number of markers available to characterize the complex microglial activation states. However, single-cell mass cytometry lacks the spatial component, which is essential when studying the relation with Aβ. Imaging mass cytometry, on the other hand, does capture the spatial distribution, however to date does not enable high-throughput analysis and offers limited resolution. Since microglia have a very complicated and variable morphology, solely evaluating protein expression directly surrounding the nucleus is insufficient, and high-resolution images are required for proper segmentation and phenotyping. Secondly, as we are studying relatively rare activated microglia subtypes that will not be present in every ROI or even subject, we required high-throughput quantitative analysis methods. The mIF-mic panel, together with our optimized microglia segmentation pipeline for 2D-images, enabled accurate segmentation and analysis of > 60000 cells to carefully identify the FTL^+^-microglia in an unbiased fashion.

In this study we adopted an unsupervised learning approach to generate distinct clusters in our dataset, and avoid bias in the identification in clusters, as can be present in more classical IHC studies. However, as already indicated in the results section, even though distinct clusters were identified, the low degree of separation on the t-SNE mapping and similarity on the associated heatmap, suggest these clusters may be more of a continuum rather that distinct subsets. This is in line with other transcriptomic and proteomic studies, in which they also showed the microglia clusters to be more of a continuum, even when studying substantially more genes or proteins[5,58,59]. However, employment of distinct clusters allows for studying the extreme ends of the continuum of the clusters to find meaningful changes in activation state. Finally, to verify that we were not looking at arbitrary differences in expression levels, we visually checked distinguishability of all independent clusters on the associated immunohistochemical images and merged clusters where this was not possible, as illustrated in supplementary Fig. 3.

Future studies looking into the effect of iron and Aβ in humanized models such as iPSC-derived microglia would be extremely valuable to decipher the functional effect of this combination, and the influence of Alzheimer-associated genetic risk variants such as APOE. In addition, since microglia, as well as iron accumulation, are shown to be involved in many different neurodegenerative and neuro-immunological disease such as Parkinson’s disease and multiple sclerosis, it would be worthwhile looking into this interaction as a common pathway in neurodegeneration. Like for Alzheimer disease, iron could interact with the accumulating protein of interest to affect microglia functioning and consequentially accelerate disease progression.

## 5. Conclusion

In summary, we showed that our multispectral immunofluorescence pipeline allowed for accurate identification of specific microglia clusters, and more importantly for the spatial analysis with respect to pathological hallmarks. In this specific study we identified dystrophic FTL^+^Iba1^+^TMEM119^-^P2RY12^-^-microglia to be significantly more present in Alzheimer’s disease patient, and to be the predominant Aβ-plaque infiltrating microglia cluster. Finally, in correspondence with the increase of FTL-expression, FTL^+^Iba1^+^-microglia showed massive iron-loading.

## Ethics approval and consent to participate

All material has been collected with written consent from the donors and the procedures have been approved by the Medical Ethical committee of the LUMC and the Amsterdam UMC.

## Consent for publication

Not applicable

## Acknowledgements

We would like to thank all patients who donated their brain to the Leiden University Medical Center (LUMC), Netherlands Brain Bank (NBB) or the Normal Aging Brain collection Amsterdam (NABCA), and prof. A.J.M. Rozemuller for neuropathological evaluation of the brains. We would also like to thank I.M. Hegeman-Klein for technical assistance with histological and immunohistochemical techniques.

## Data availability

The data that support the findings of this study are available from the corresponding author upon reasonable request.

## Authors’ contributions

B.K. and L.v.d.W. conceived and designed the project. B.K., M.I. and N.F.C.C.d.M designed the antibody panel for microglia multispectral immunofluorescence (mic-mIF). A.S., O.D. and B.K. created the microglia segmentation pipeline. A.S. created the spatial analysis tools for mic-mIF data under supervision of B.P.F.L, J.D. and T.H.. B.K and L.d.H. performed morphological evaluation of microglia. B.K. and A.S., analysed and interpreted the mic-mIF data. B.K., A.S., W.M.C.v.R-M, T.H., and L.v.d.W. wrote the manuscript. All authors read and approved the final manuscript.

