## Supplementary files for "Iron-loading is a prominent feature of activated microglia in Alzheimer’s disease patients"

### Supplementary Material

**Supplementary Table 1**. **Patient demographics.** ** Unknown pathology score. DO = Disease onset. PM = Post-mortem. LOAD = Late onset Alzheimer's disease. EOAD = Early onset Alzheimer's disease

| | **Diagnosis** | **Sex** | **DO** | **Age** | **Braak** | **Amyloid load** | **PM delay** | **APOE genotype** | **Familial genetic variant** |
|---|---|---|---|---|---|---|---|---|---|
| C1 | Control | F | | 91 | 2 | 2/3 | 03:47 | 3/3 | |
| C2 | Control | M | | 73 | 2 | 1 | 08:00 | 3/3 | |
| C3 | Control | M | | 82 | ** | ** | 05:30 | | |
| C4 | Control | F | | 87 | 1 | 0 | 08:30 | 3/3 | |
| C5 | Control | F | | 72 | 0 | 0 | 07:15 | 3/3 | |
| C6 | Control | M | | 93 | ** | ** | 08:30 | | |
| C7 | Control | M | | 82 | 1/2 | 1/2 | 07:30 | 3/4 | |
| C8 | Control | F | | 89 | 3 | 1/2 | 06:30 | 3/4 | |
| C9 | Control | F | | 72 | 1 | 1 | 06:50 | 3/3 | |
| AD1 | LOAD | F | 80 | 88 | 6 | 5 | 04:40 | 3/4 | |
| AD2 | LOAD | M | 69 | 73 | 5 | 5 | 04:45 | 4/4 | |
| AD3 | LOAD | F | >65 | 82 | 4 | 4 | 04:35 | 3/4 | |
| AD4 | EOAD | F | 60 | 73 | 6 | 5 | 07:17 | 3/3 | |
| AD5 | EOAD | M | 61 | 72 | 6 | 5 | 05:15 | 3/4 | |
| AD6 | EOAD | F | 40 | 67 | 6 | 4 | 04:30 | 3/4 | |
| AD7 | EOAD | F | 64 | 91 | 6 | 4/5 | 04:20 | 3/3 | |
| AD8 | EOAD | M | 47 | 59 | 5 | 4/5 | 05:25 | 3/3 | APP duplication |
| AD9 | LOAD | F | 85 | 90 | 6 | 4/5 | 03:55 | 2/3 | |
| AD10 | EOAD | F | 51 | 70 | 6 | 5 | 04:20 | 4/4 | |
| AD11 | LOAD | F | 87 | 89 | 4 | 3-4 | 04:30 | 3/3 | |
| AD12 | EOAD | F | 34 | 43 | 6 | 5 | 04:15 | 3/3 | PSEN1 |

**Supplementary Table 2. Details microglia multispectral immunofluorescence (mic-mIF) panel**

| | **Staining target** | **Antibody** | **Isotype** | **Antigen retrieval** | **dilution** | **Incubation time** | **Secondary Conjugate** |
|---|---|---|---|---|---|---|---|
| 1. | P2RY12 | HPA014518, Sigma Aldrich | rIgG | 10mM Citrate buffer (pH=6.0) | 1:2500 | 2h | Poly HRP + Opal 520 |
| 2. | TMEM119 | HPA051870, Sigma Aldrich | rIgG | | 1:250 | ON | Poly HRP + Opal 570 |
| 3. | Light Chain Ferritin (FTL) | AB69090, Abcam | rIgG | | 1:100 | ON | G-α-Rab Alexa 594 |
| 4. | Aβ (17–24) | SIG-39220, Biolegend | mIgG2b | | 1:250 | ON | Goat-α-mIgG2b Alexa 647 |
| 5. | Iba1 | MABN92, Millipore | mIgG1 | | 1:20 | ON | G-α-mIgG1 CF680 |
| 6. | Nucleus, DAPI | D9542-1mg, Sigma Aldrich | n.a. | | 0.1 µg/ml | 5 min | n.a. |

**Supplementary Table 3. Key resources**

| Product | Catalogue nr. | Supplier | Dilution |
|---|---|---|---|
| **Primary antibodies** | | | |
| Purified anti-β-Amyloid | SIG-39220 | Bio legend | 1:250 |
| Anti-Iba1/AIF1 | MABN92 | Millipore | 1:20 |
| Anti-Ferritin Light Chain (rab) | ab69090 | Abcam | 1:100 |
| Anti-Ferritin Light Chain (mIgG2a) | SC-74513 | Santa Cruz | 1:100 |
| Anti-P2RY12 | HPA014518 | Sigma Aldrich | 1:2500 |
| Anti-TMEM119 | HPA051870 | Sigma Aldrich | 1:250 |
| **Secondary antibodies** | | | |
| Opal 520 Reagent Pack | FP1487001KT | Perkin Elmer | 1:100 |
| Opal 570 Reagent Pack | FP1488001KT | Perkin Elmer | 1:100 |
| Goat anti-Rabbit IgG (H+L), Alexa Fluor 594 | A-11037 | ThermoFisher | 1:200 |
| Goat anti-Mouse IgG2b, Alexa Fluor 647 | A-21242 | ThermoFisher | 1:200 |
| Goat anti-mouse IgG1, CF680 | 20253 | Biotium | 1:200 |
| Goat anti-Mouse IgG (H+L), Alexa Fluor 546 | A-11003 | ThermoFisher | 1:200 |
| Goat anti-Rabbit IgG (H+L), Alexa Fluor 546 | A-11010 | ThermoFisher | 1:200 |
| Swine Anti-Rabbit Immunoglobulins/Biotin | E0353 | DAKO | 1:400 |
| Rabbit anti-Mouse Immunoglobulins/Biotin | E0354 | DAKO | 1:200 |
| VECTASTAIN Elite ABC-HRP Kit, Peroxidase | PK-6100 | Vector Laboraties | n.a. |
| **Extra resources** | | | |
| BrightVision+ Poly- HRP-Anti Mouse/Rabbit IgG Biotin-free | VWRKDPVO1 10HRP | Immunologic | n.a. |
| 1X Plus Amplificiation Diluent | FP1498A | Perkin Elmer | n.a. |
| ProLong Diamond Antifade mountant | P36961 | ThermoFisher | n.a. |
| DAPI | D9542-1mg | Sigma Aldrich | 0.1ug/mL |
| DAB sigma | D5637 | Sigma Aldrich | 0.5ug/mL |

**Supplementary Fig. 1**. Sample images of DAB-enhanced, single immunofluorescence and multiplexed immunofluorescence stainings of all antibodies used in our multispectral immunofluorescence panel. Scale bar, 100 μm.

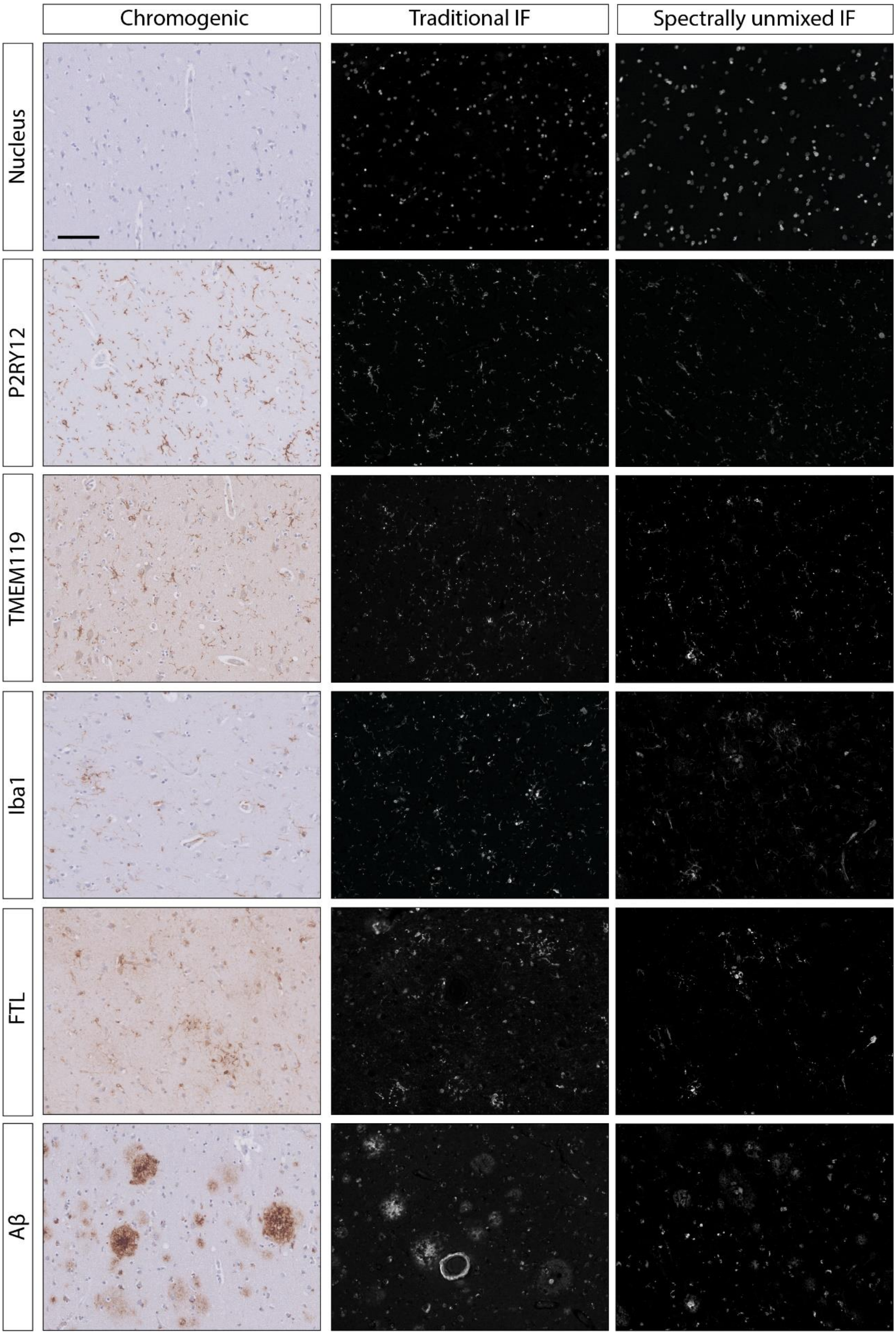

**Supplementary Fig. 2.** Example of multispectral immunofluorescence data extracted from the grey matter of an AD-patient**.** The segmentation mask derived: manually by the specialist (**b**), using the GliaTrace toolbox (**c**), automatically from the "Inform" image acquisition software (**d**) and from our proposed segmentation pipeline (**e**). **f** Dice's coefficient for each of the 156 cells segmented by our method. Number of correctly identified (**g**), missing (**h**) and falsely identified cells (**i**).

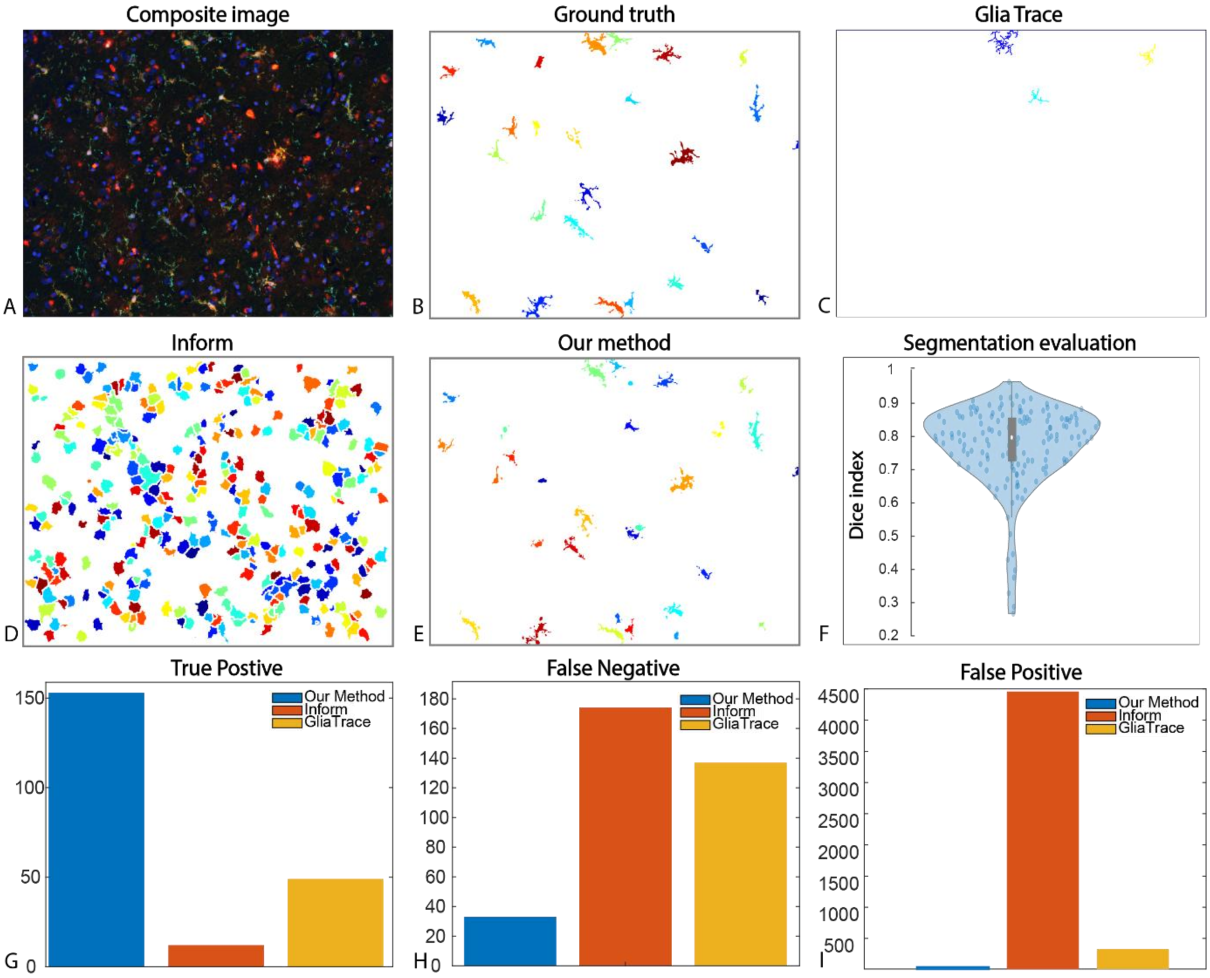

**Supplementary Fig. 3.** Sample images of all original clusters, and how they were merged. Scale bar, 20 μm.

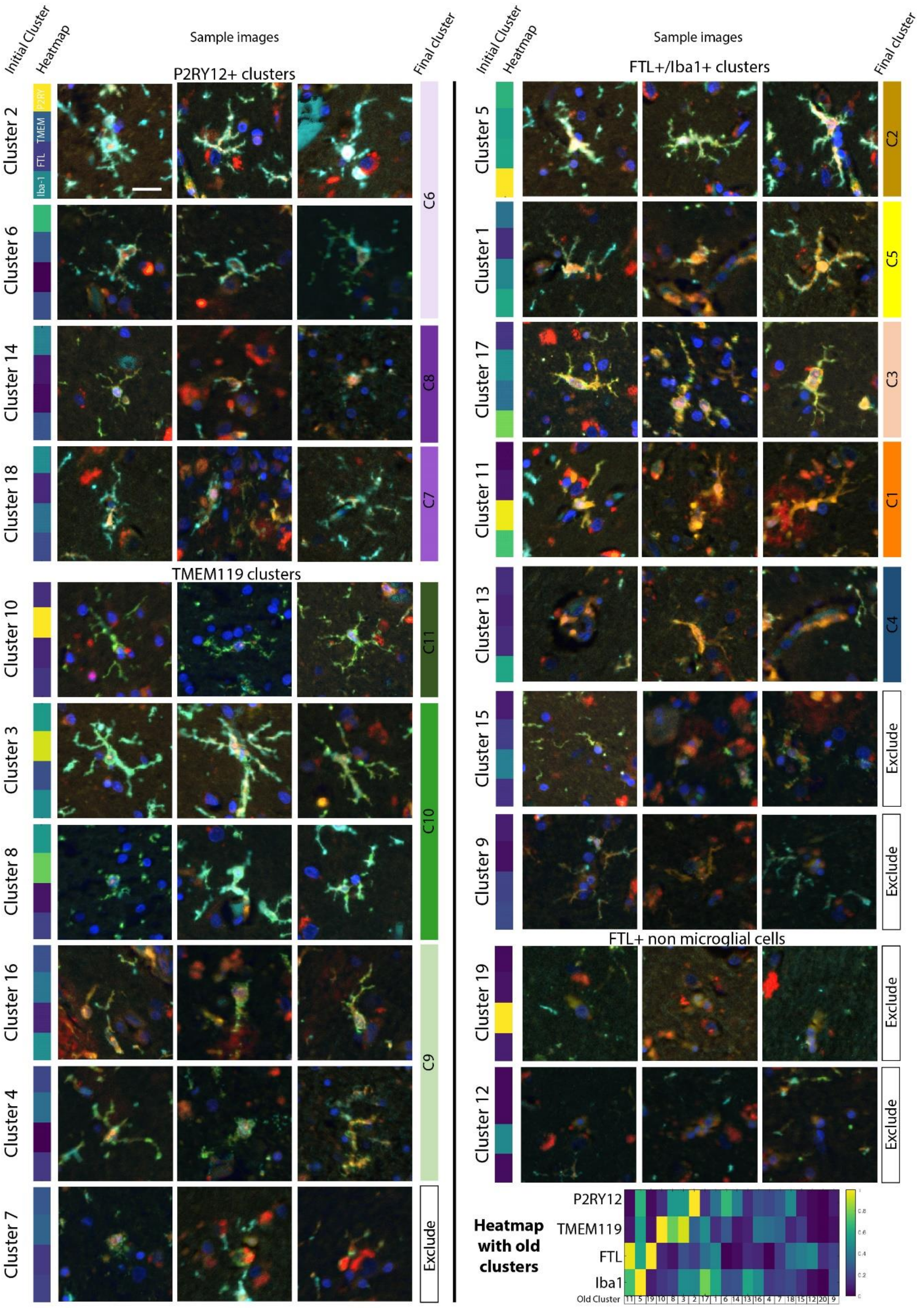

**Supplementary Fig. 4.** Correlation of iron$^{+}$-microglia with C2-microglia (**a**) and C3-microglia (**d**) is nonsignificant ($n$ = 20, Pearson coefficient). C2 microglia are mostly present in controls with early stage tau and amyloid-β pathology (**b-c**), whereas C3 microglia are variably present among all stages in both controls and Alzheimer patients (**e-f**). Comparison between LOAD ($n$ = 5) and EOAD ($n$ = 7) patients shows no differences in number of identified Aβ-plaques (g), percentage of microglia infiltration (**e**), prevalence of all-mic (**i-k**), prevalence of Aβ-mic (**l-n**), and percentage of microglia infiltrating Aβ-plaques (**o-q**) for C1, C2 and C3-microglia (Median, Mann-Whitney U test). Comparison between APOE4 ($n$ = 6) vs. APOE3 ($n$ = 4) shows no difference in prevalence in all-mic(**r,s**), nor Aβ-mic (**t,u**) of C2 and C3 microglia. A significantly increased proportion of C2-microglia infiltrated Aβ-plaques (v), but this was not the case for C3-microglia (**w**) (Median, Mann-Whitney U test).

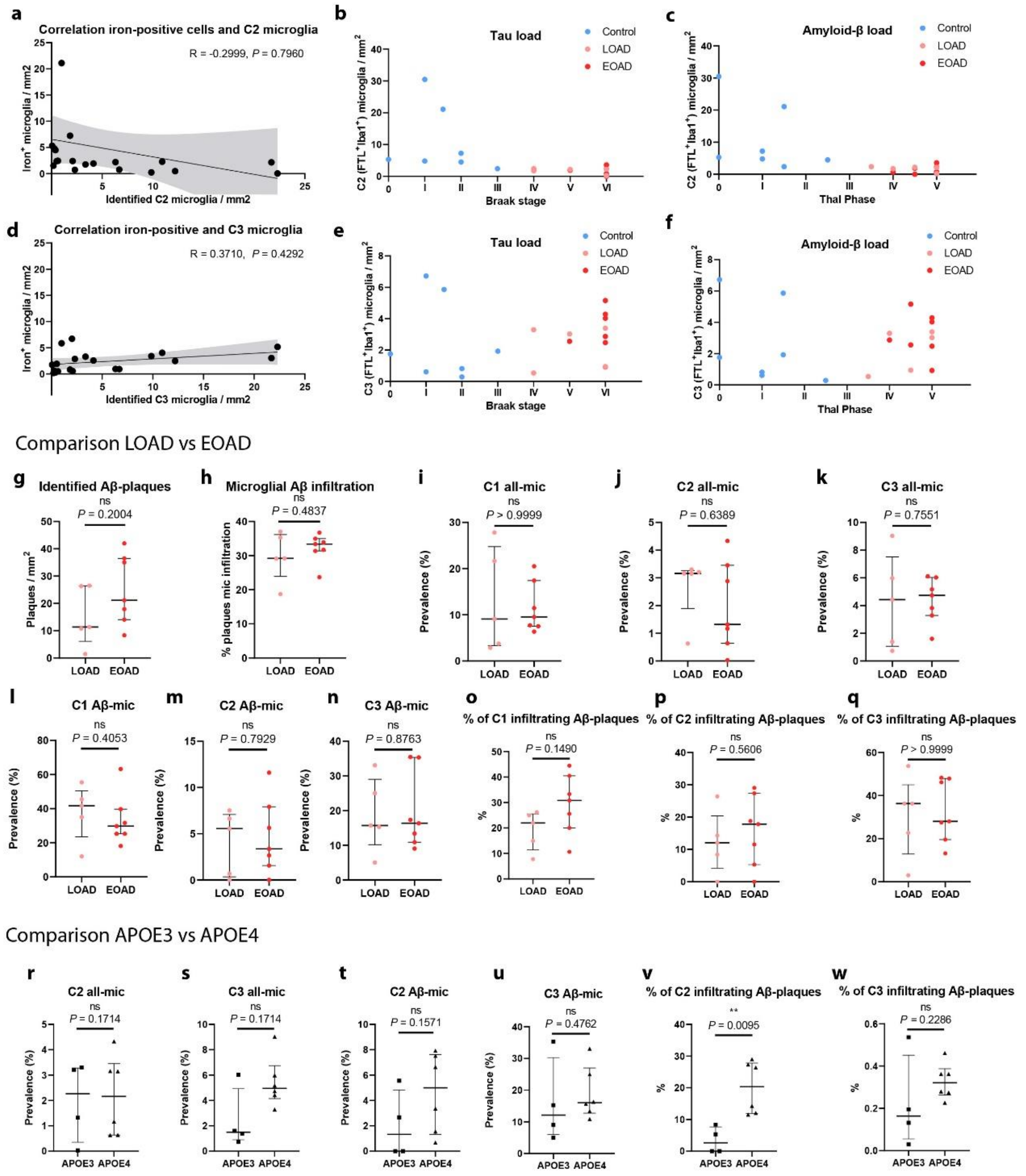

**Supplementary methods**

**1. Histological staining protocol**

10-μm-thick sections were used for histochemical iron detection using an enhanced Perl's reaction previously published by Van Duijn *et al.* (2013). Sections were incubated for 80 min in 1% potassium ferrocyanide, washed three times in 0.1 M phosphate buffer, followed by 100 min incubation in methanol with 0.01 M $NaN_3$ and 0.3% $H_2O_2$. Subsequently, sections were washed again and incubated for 80 min in a solution containing 0.025% 3′3-diaminobenzidine-tetrahydrochloride (DAB, Sigma) and 0.005% $H_2O_2$ in 0.1 M phosphate buffer. The reaction was stopped by washing with tap water. A consecutive 10-μm-thick section was used for IHC detection of FTL. Sections were deparaffinized with xylene, washed with alcohol and blocked with 0.3% $H_2O_2$/methanol for 20 min. Subsequently, they were incubated with FTL (1:100, Santa Cruz) overnight at room temperature and after washing with PBS incubated with anti-rabbit alkaline phosphatase (1:50, Vector) for 1 h. Sections were washed with PBS and incubated with Vector Blue for 20 min in the dark. Finally, they were rinsed and covered with aqua mount.

**2. Brightfield microscopy**

Chromogenically stained slides were imaged using a Philips IntelliSite Ultra Fast Scanner (Philips, the Netherlands ). Whole slide images could be viewed in 2×–40× magnification using the Philips Intellisite digital Pathology Solution system. Whole slide images were exported at 4× magnification for quantification of number of iron-positive cells. Snapshots of higher magnification were taken directly in the image-viewer.

**3. Antibody validation for multiplexed IHC**

The six-colour mic mIF panel (Supplementary Table 2) was created and optimized following a previously described protocol (IJsselsteijn *et al.*, 2019). Individual antibody conditions were optimized using single IHC and IF. Firstly, individual antibodies were tested with chromogenic and fluorescent detection for optimal antibody concentration and antigen retrieval method with the following protocol. 5-μm-thick FFPE tissue sections were deparaffinized with xylene, washed with alcohol and endogenous peroxidase was blocked by incubating in 0.3% H202 in methanol for 20 min. Antigen retrieval was performed with either citrate buffer (10 mM, pH = 6.0) or EDTA buffer (10 mM, pH 9.0). After cooling, the sections were blocked with 0.1% BSA/PBS with 0.05% Tween for 30 min and incubated with primary antibody diluted in blocking buffer in a range of concentrations overnight at room temperature. For chromogenic detection, the slides were washed with PBS and incubated with Sw-a-Rb/biotin (1:400, Dako) or Sw-a-Mouse/biotin (1:200, DAKO) for 60 min, followed by 30 min incubation with VECTAstain elite ABC. Chromogenic substrate was developed with 0.05% DAB (Sigma) with 0.005% $H_2O_2$ for 10 min. The reaction was stopped with tap water, after which the sections were counterstained with haematoxylin for 5 min and mounted with micromount. For fluorescent detection, slides were incubated with the appropriate Alexa 546 fluorophore (1:200, ThermoFisher) for 1 hour. Subsequently, the slides were incubated with 0.1 μg/mL DAPI (Sigma) for 5 min and mounted with prolong diamond (ThermoFisher). After individual antibody testing, antibodies were combined to test the viability for multiplexed immunofluorescence. The full protocol described in section 2.3 was performed for 7 slides of one control and one AD subject. One slide of each was incubated with all primary antibodies. For each of the other slides the exact same protocol was performed, but only 1 out of the 6 primary antibodies was added in the primary incubation step. These slides were used to analyse the fluorescent spectrum of the individual fluorophores on our slides, which are subsequently used to un-mix the 6 different spectra. To verify specificity, the extracted signal was compared with single immunofluorescence (Supplementary Fig. 1).

**4. Multispectral microscopy and image-acquisition**

Mic-mIF-stained tissue slides were scanned at 4× magnification using the Vectra 3.0 Automated Quantitative Pathology Imaging system (PerkinElmer). Following whole slide scanning, a 50% and 25% ROI grid was placed on the cortex and white matter, respectively. For each ROI, high-resolution 20× magnification images are obtained of all subjects. Spectral separation of the 6 individual dyes was performed automatically using InForm Cell Analysis software (PerkinElmer), using spectral libraries obtained with single-marker IF detection of the different fluorophores. In total, six different raw component images of the extracted spectra were exported from Inform for further analysis.

**5. Confocal microscopy**

20 μm sections were imaged using an Andor Dragonfly 200 spinning disk confocal system (Andor, Oxford Instruments). Sections were stained with DAPI, Alexa 488, Alexa 546 and Alexa 647, which were imaged with a 405 nm, 488 nm, 561 nm and 637 nm laser respectively. Subsequently images were exported and processed using Imaris (Bitplane, Oxford instruments). First, a Gaussian filter was applied to all channels, after which snapshots of the reconstructed 3D projection were taken.
